# Automatic measurement of temporalis muscle thickness from CT head scans using deep learning

**DOI:** 10.1101/2025.10.31.685721

**Authors:** Nicole Hernandez, Tommi K. Korhonen, Emilia K. Pesonen, Sami Tetri, Lasse Pikkarainen, Said Pertuz, Otso Arponen

## Abstract

**Objective:** To construct and validate a deep-learning (DL) model for the automatic quantification of temporalis muscle thickness (TMT) in CT head scans.

**Materials and methods:** We developed and evaluated the performance of a DL-based method for the measurement of temporalis muscle thickness using publicly available CT head scans. Reference standard TMT was established using a previously published measurement protocol, applied on 198 CT head scans obtained from a publicly available database, originally collected from various radiology centers within one city in 2017. A DL landmark detection model was trained to measure the temporalis muscle thickness. The absolute error and correlation between DL-based and reference standard TMT measurements were calculated. Additionally, the ability of the DL-based measurements to stratify subjects into low TMT vs normal TMT groups was assessed using the metrics of specificity, sensitivity, and accuracy.

**Results:** The median reference TMT value was 6.0 mm (95% CI: 5.4, 6.5); the median DL-based TMT value was 5.8 mm (95% CI: 5.6, 6.3). The mean absolute error for TMT was 0.7 mm (95% CI: 0.6, 0.9). The correlation coefficient between reference and DL-based TMT was 0.9 (95% CI: 0.8, 0.9). The DL-based measurements classified the patients into low and normal TMT groups with sensitivity of 84.2%, specificity of 85.0% and accuracy of 84.6%.

**Conclusions:** Our DL-based pipeline allows for fully automated and reproducible quantification of temporal muscle thickness and patient stratification into low and normal TMT groups.

**Summary statement:** In this study we developed and tested a deep learning model for the measurement of temporal muscle thickness in CT head scans, showing potential utility for clinical decision making.

**Key points:**

- Temporalis muscle thickness surrogates muscle mass, which is associated with frailty; however, there is no automatic tool to measure temporalis muscle thickness in CT head scans
- Our deep learning model-based temporalis muscle thickness measurements presented a strong, positive linear correlation with reference standard measurements, and less than 1mm mean absolute error
- The deep learning-based measurements demonstrated strong potential for stratifying subjects into low versus normal muscle levels, enabling quick assessment of temporalis muscle status.

## Introduction

Loss of skeletal muscle mass and strength are a key feature in several conditions associated with increased vulnerability, such as sarcopenia, cachexia, malnutrition, and frailty (1). Low muscle mass, most commonly measured at the level of the third lumbar vertebra (L3), has been identified as a radiological marker of loss of skeletal muscle mass and impaired survival in several diseases (2). However, patients with neurological conditions often only undergo imaging of the head. The opportunistic use of these clinical scans could enable timely evaluation of muscle mass without additional radiation exposure or costs. Temporal muscle thickness (TMT) surrogates skeletal muscle mass, is associated with L3 muscle area (3,4), grip strength (3,5,6), and has shown prognostic value in patients with chronic subdural hematoma (7,8), stroke (9,10), and different types of cancer (11,12).

Manual assessment of TMT requires human resources, prior training, and is subject to inter-reader variability, highlighting the need for automated measurement workflows. Previously, TMT has been measured from MRI images using deep learning (DL) (13–15), but automated measurement of TMT from CT data has not been demonstrated to date. CT is more widely available than MRI, and often the only imaging modality used in the acute phase in many neurological diseases (16). Developing automated workflows for the measurement of TMT from CT would enable prompt recognition of patients with low muscle mass, to aid in clinical decision-making and directing ancillary care to patients likely benefiting from it.

To this end, we constructed a DL model to measure the temporalis muscle thickness from CT slices according to a previously published, validated measurement protocol (8,17). We then assessed the utility of the DL model in stratifying subjects into low and normal TMT groups.

## Methods

### Study design

This study exclusively involved anonymized imaging data retrieved from an open online repository, and no personal, identifiable information was analyzed nor retrievable; accordingly, institutional review board approval and patient informed consent were not required. This is a study on the development and evaluation of a DL-based method for the measurement of temporalis muscle thickness using publicly available CT head scans.

### Data

The data used in this study was a subset of the CQ500 dataset (18). Briefly, CQ500 includes 491 anonymized DICOM-format CT head scans of patients diagnosed with head or brain injuries. The mean age of the patients in the cohort is 48.1 years (SD: 21.6). Information on the patients’ sex, age, acquisition machine, and site was not made available with the release.

The test sample size was estimated to achieve a 95% confidence interval for the mean absolute error between DL-based and reference standard TMT with a half-width of 0.2 mm. The half-width value was chosen to ensure high precision of the error estimation, while considering practical feasibility given available data. A standard deviation of 0.6 mm for the mean absolute error based on (14) was used for calculations. The calculated sample size was 39 subjects. Based on this, we selected a total of 198 scans for the total dataset, including training and validation data. We only included non-contrast scans of thickness and spacing smaller than 0.5 mm.

To reduce spatial variability between subjects, all 198 scans were manually registered, and the axial level at which to measure the TMT was defined. To this end, two clinicians, E.K.P. with 4 years of experience in clinical imaging research, and O.A. with 10 years of experience in clinical imaging research), each processing 50% of the data, manually registered the scans and defined the TMT measurement level on the orbital roof. The scans were converted into the nii.gz format. The slice at the measurement level was extracted and resampled to pixel size 0.5 mm. To simplify the model task and avoid training separate models for each side of the head, the images were automatically divided into left and right hemispheres by calculating a midline that divided the skull area into equal left and right sides, the hemispheres were separated into individual images, and the left hemisphere was mirrored to resemble the orientation of the right hemisphere. To exclude irrelevant regions, the skull area was identified via thresholding, and used to remove the skull, brain and pixels that were farther than 2cm from the skull. Lastly, the images were thresholded to values between 0 and 80 HU (see Figure 1) and resized to a standard size of 512 x 256 pixels.

**Figure 1.**
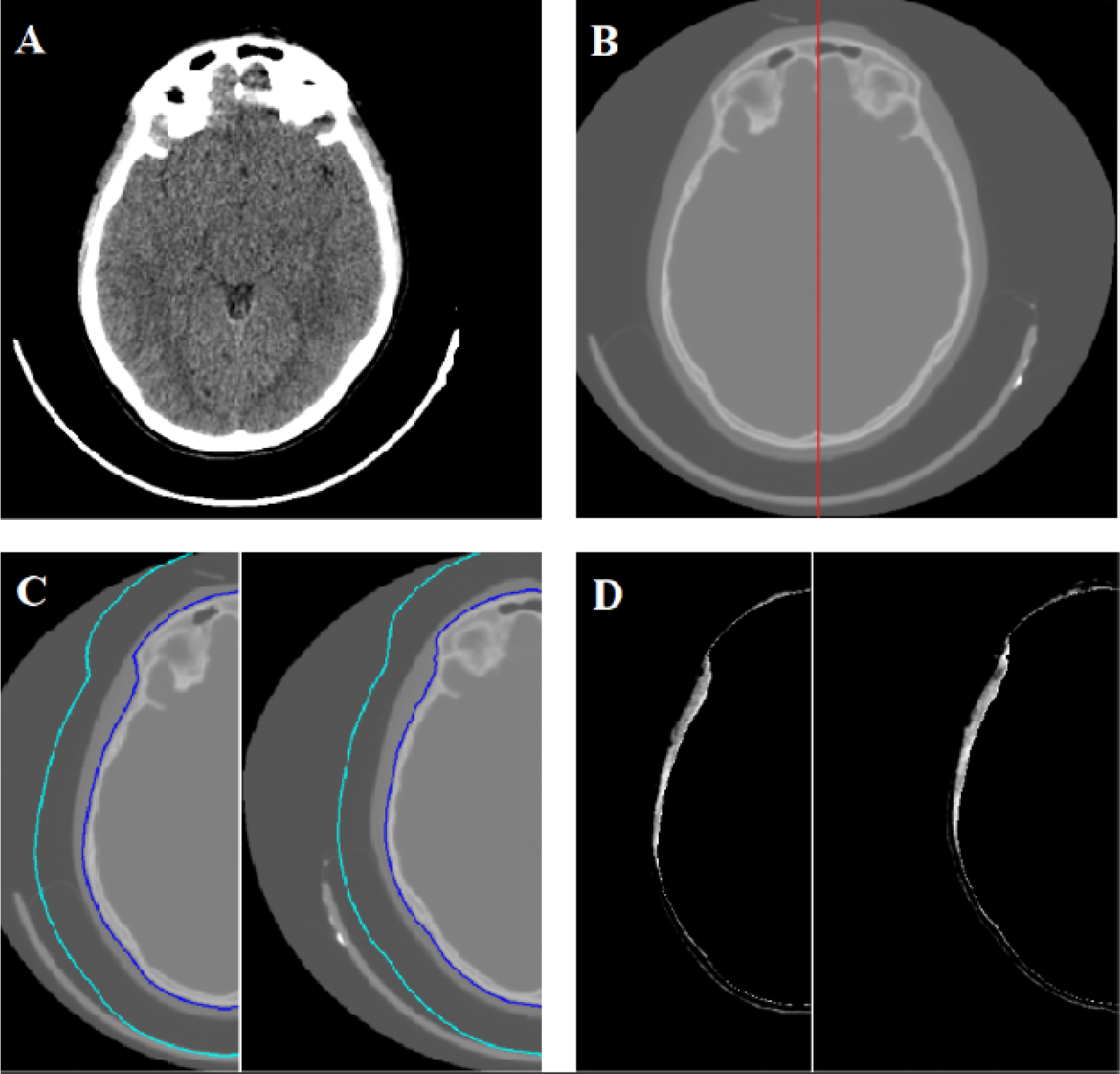
Scan preprocessing. A: Original scan windowed to linear width/level 80/40 Hounsfield units, used by the expert clinician to define TMT. B: The original slice was resampled to pixel size of 0.5 mm and the midline (red line) that divides the skull area into equal halves was defined. C: The middle line was used to divide the slice into two images; the left side was mirrored horizontally. The exterior edge of the skull (blue line) was used to define a line 2 cm away (cyan line). D: The area encompassed between the blue and cyan lines was preserved and the area outside it set as background; the image was thresholded to Hounsfield units between 0 and 80. Abbreviations. TMT: temporalis muscle thickness.

### Reference standard

Reference standard TMT was measured in scans windowed to linear width/level 80/40 Hounsfield units (HU) from the thickest part of the temporal muscle perpendicular to the line connecting the external tables of the most medial part of the temporal fossa and the most lateral part of the temporoparietal bone, according to a previously validated protocol (8,17). The reference standard was established by E.K.P., who defined the endpoints of the line segments denoting the thickness of the left and right temporal muscles in all 198 scans, and exported them in the.JSON format. These procedures were conducted in 3D Slicer v.5.0.3.

### Data partitions

A random train/tuning/test partition of 120 (∼60%) /39 (∼20%) /39 (∼20%) was used. Each partition was derived from the full group consisting of the 198 scans included in the study.

### Model

We developed a two-stage TMT measurement model. The first stage is aimed at the automatic detection of landmark points for measuring TMT. As illustrated in Figure 2, two external points (E1 and E2) and two internal points (I1 and I2) are used to draw four candidate TMT measurements. For the detection of these points, we trained a landmark detection model, the PyTorch-based Python package Landmarker, a Toolkit for Anatomical Landmark Localization in 2D/3D Images (19). Specifically, the Landmarker implementation of the Spatial Configuration Network (SCN) (20). The SCN network was created for the localization of landmarks in medical image applications, in scenarios with limited amounts of training data. The SCN reduces the need for large amounts of data by dividing the landmark localization problem into two simpler problems that are, relatively, less data hungry: heatmap regression using a convolutional neural network, and spatial configuration knowledge using consecutive convolutional layers.

**Figure 2.**
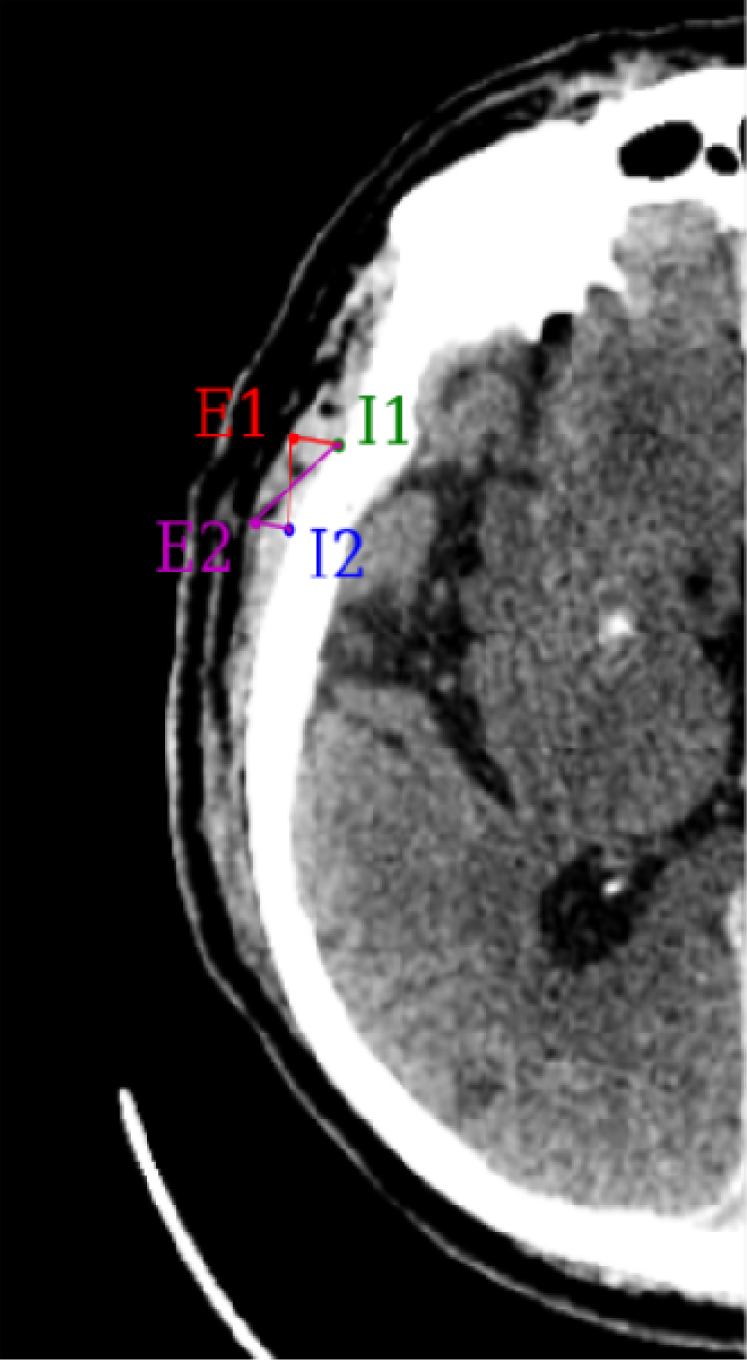
Four coordinates were generated using the trained landmark detection model: two possible coordinates for the exterior landmark (E1 and E2) of the temporalis muscle, and two for the interior (I1 and I2). The four possibilities for the outcome are represented by the possible segments: ————. The pair of landmarks with the highest similarity to the tuning dataset was selected.

The second stage of our model receives the four candidate measures and selects one using nearest neighbors interpolation from training data. Any given candidate corresponds to a landmark pair denoted by the extreme points *E* and *I* (see Figure 2), of coordinates (*x*_*E*_, *y*_*E*_) and (*x*_*I*_, *y*_*I*_), respectively. Each candidate was represented by a vector of six features consisting of: the four coordinates that define the landmark pair, *x*_*E*_, *y*_*E*_, *x*_*I*_, *y*_*I*_; the Euclidian distance between the landmarks, i.e., the norm of the segment, 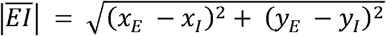; and the angle ϑ between the segment 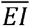 and the vertical axis, 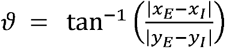. The distance of each candidate vector to the six-feature vector set generated from the training reference values was calculated, and the DL-based landmark pair corresponding to the vector with the smaller distance was chosen. The DL-based TMT was obtained by calculating the Euclidean distance between the chosen DL-based landmarks. The full pipeline is presented in Figure 3.

**Figure 3.**
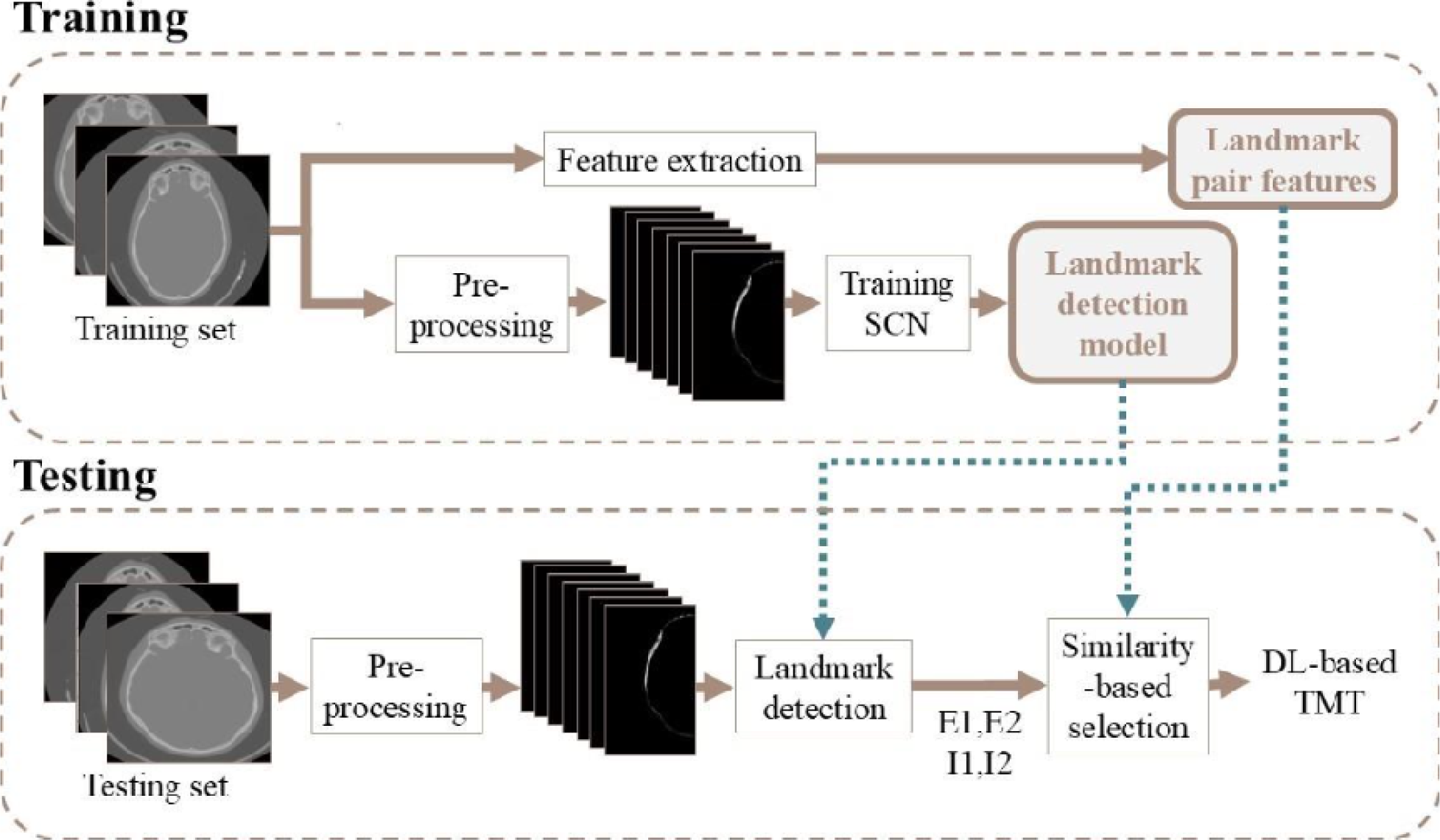
Automatic TMT assessment pipeline. In the training phase, the CT scans and respective landmarks were preprocessed and used to train a landmark detection model. During the testing phase, the landmark detection model was used to generate 4 possible pairs of landmarks per temporalis muscle, and similarity-based detection was used to select the most likely pair that represents the temporalis muscle thickness. Abbreviations. DL: deep learning, TMT: temporalis muscle thickness.

All experiments were conducted in a computing environment equipped with two Intel Xeon processors, with 20 cores each running at 2.1 GHz and one Nvidia Volta V100 GPU. The packages used were Optuna (4.3.0), NiBabel (5.3.2), MONAI (1.4.0), Landmarker (0.2.1), SimpleITK (2.5.0), MLflow (3.1.1), NumPy (1.26.4), pandas (2.3.0), tqdm (4.67.1), PyTorch (2.5.2), CUDA (12.9.0) and Python (3.11).

### Training

Transfer learning and data augmentation were used during the training phase, due to the limited dataset size. Weights obtained from training the SCN (20) for the localization of 37 landmarks in hand radiographs were retrieved. The retrieved weights were used to initialize the most internal down sampling layers of our architecture. All the layers of our model were trainable.

We augmented our training dataset three-fold using the MONAI framework (21) and implemented transformations of the following type: rotations between ±15°, translation of ±10 pixels, scaling of zoom ±10%, shearing of ±0.1, random contrast adjustment of gamma between 0.5 and 4.5 – gamma correction adjusts contrast via a power-law transformation –, gaussian noise with mean 0 and standard deviation of 0.1, random intensity scale by up to 25% and random histogram shift, these last three with a low probability of 20%, and intensity normalization between zero and one. The random nature of these transformations ensured that no two images were the same in the augmented training data.

Optuna (22) was used for hyperparameter exploration, and the training loss, tuning loss, and mean point error (MPE) on the tuning set were monitored using MLflow. The point error is the Euclidian distance between the reference standard and DL-based landmarks, which is then averaged over all landmarks and images. We used Optuna’s TPEsampler sampling algorithm and pruning to skip unpromising trials.

The top three performing models were selected by identifying the minimum tuning MPE. Their performance was then evaluated in the testing set and the model with the lowest test MPE was selected for analysis.

### Evaluation

The performance of our pipeline was evaluated on the test set. DL-based TMT was assessed using the absolute and relative errors, correlation coefficient, and statistical differences using a Wilcoxon signed-rank test for differences at the confidence level of 5%. We provided a fuller picture of the absolute and relative errors by reporting mean, median, and confidence interval values. Additionally, the DL-based landmarks were assessed using the MPE.

Studies on the utility of TMT to assess muscle tone define a TMT cut-off value to stratify patients into low or normal muscle mass groups (23). A cut-off value for normal muscle mass has not been agreed upon in the literature, and the values used in each study have been highly dataset-dependent: numerous studies used the median of their TMT distribution as the cut-off value (12,15). Similarly, we used the median of the reference values distribution as the cut-off between low and normal TMT. In this scenario, the capacity of the DL-based TMT measurements to accurately classify subjects into low TMT and normal TMT groups was assessed using the metrics of sensitivity, specificity, and accuracy. It is important to note that, in numerous studies, the TMT of a specific subject corresponds to the average of their left and right TMT values, and the median TMT value used as cut-off corresponds to the median value of the average left-right TMT distribution. For this section of the analysis, we replicated these calculations and presented results at the subject level, even though we keep using the simpler terms of TMT and median TMT for readability. All analyses were performed in Python.

## Results

### Performance metrics and diagnostic performance metrics

The total MPE for landmark localization was 5.2 mm in the tuning set and 7.5 mm in the test set.

The median values and confidence intervals for the reference standard and DL-based TMT were 6.0 mm (95% CI: 5.4–6.5) and 5.8 mm (95% CI: 5.6–6.3), respectively (see Figure 4). We found no statistically significant difference when comparing reference standard vs DL-based TMT measurements (*p* = 0.05).

**Figure 4.**
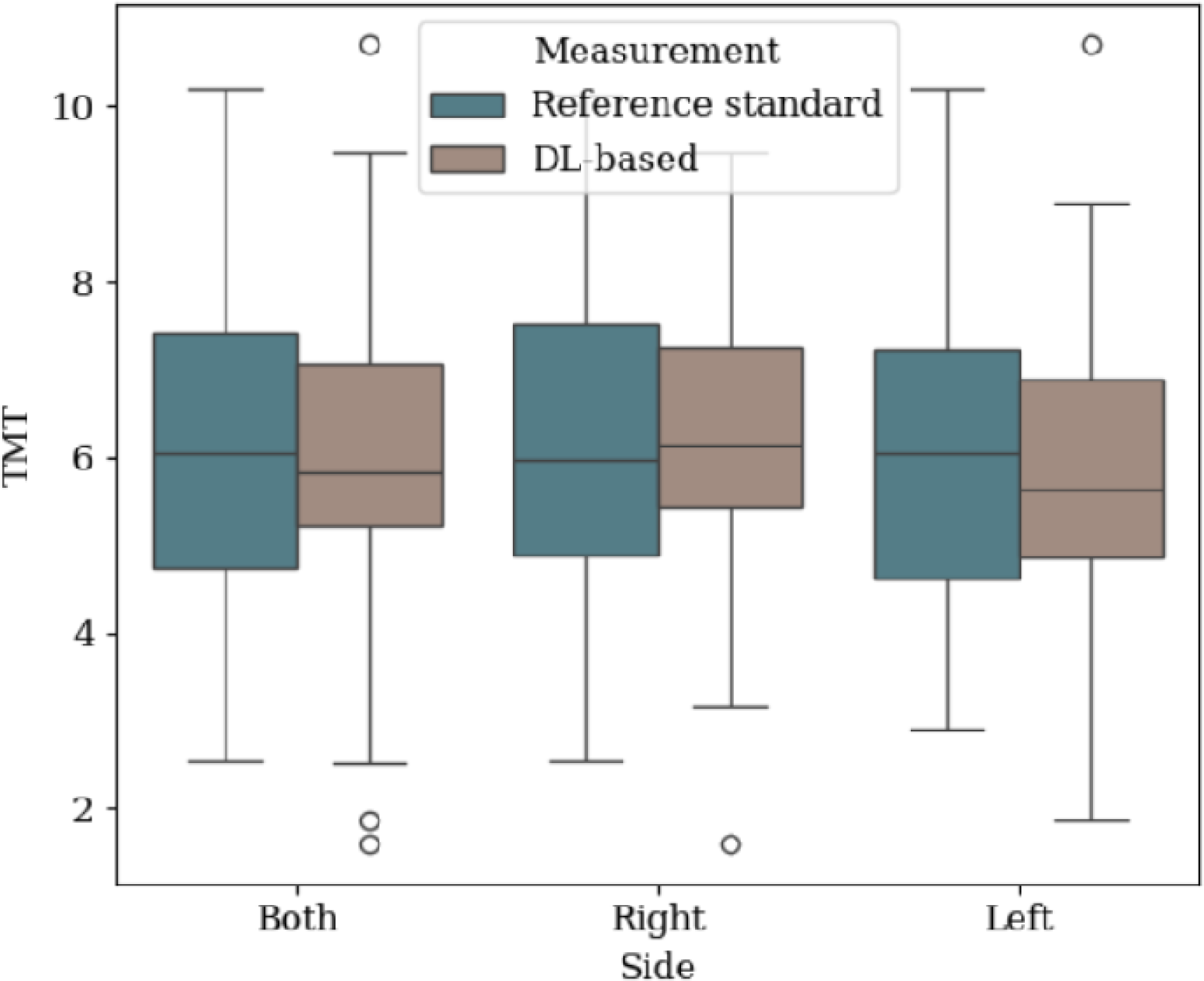
Box-plots of the distribution of reference standards and DL-based TMT, for all measurements regardless of laterality, right measurements only, and left measurements only. Abbreviations. DL: deep learning, TMT: temporalis muscle thickness.

The mean and median absolute errors were 0.7 mm (95% CI: 0.6–0.8) and 0.6 mm (95% CI: 0.5– 0.7), respectively. The mean and median relative error were 13.3% (95% CI: 10.5–16.2) and 8.8% (95% CI: 7.3–12.0), respectively. Details for all measurements combined, as well as separately for the left and right sides can be found in Table 1.

**Table 1.** Errors of the TMT estimations. Mean of the absolute and relative errors between the reference standard TMT and the DL-based TMT, measured in mm.

| Side | Absolute error [mm] (95% CI) | Relative error [%] (95% CI) |
| --- | --- | --- |
| **Both** |  |  |
| Mean | 0.7 (0.6–0.8) | 13.3 (10.5–16.2) |
| Median | 0.6 (0.5–0.7) | 8.8 (7.3–12.0) |
| **Right** |  |  |
| Mean | 0.8 (0.6–1.0) | 15.6 (10.8–20.3) |
| Median | 0.7 (0.5–1.0) | 10.9 (8.4–18.3) |
| **Left** |  |  |
| Mean | 0.6 (0.5–0.8) | 11.3 (8.2–14.4) |
| Median | 0.5 (0.3–0.9) | 8.2 (5.7–12.4) |

Abbreviations. TMT: temporal muscle thickness; DL: deep learning; CI: confidence interval.

The Pearson’s correlation coefficient between DL-based and reference standard TMT were 0.9 (95% CI: 0.8–0.9, *p* < 0.05) for all measurements as well as only right measurements, and 0.9 (95% CI: 0.9–1.0, *p* < 0.05)) for only left measurements.

In terms of stratification into low and normal TMT groups, the sensitivity, specificity, and accuracy were 84.2%, 85.0%, and 84.6%.

### Failure analysis

We calculated the Spearman rank-order correlation coefficient to test whether MPE and absolute error are positively associated. The obtained coefficient was 0.2, indicating a positive yet weak correlation between MPE and absolute error.

In the stratification scenario, six (15.4%) DL-based TMT measurements resulted in incorrect classification: three false positive (subjects with normal TMT classified as low TMT) and three false negatives (subjects with low TMT classified as normal TMT). These cases are shown in Figure 5. We looked closer at the MPE and absolute error of the incorrect classifications – false positives and false negatives – and compared them to the errors of the correct classifications – true positive and true negatives –. The median MPE of the correct classifications was 2.57mm (IQR: 1.5—9.3); the MPE of the incorrect classifications were in the range of 5.0—16.4 mm, and 50% of these measurements fell below the upper bound of the interquartile range of the correctly classified group. The median absolute error of the correct classifications was 0.6 mm (IQR: 0.4—0.9); the absolute errors of the incorrect classifications were in the range of 0.4—1.0 mm, and 83% of these measurements fell below the upper bound of the interquartile range of the correctly classified group.

**Figure 5.**
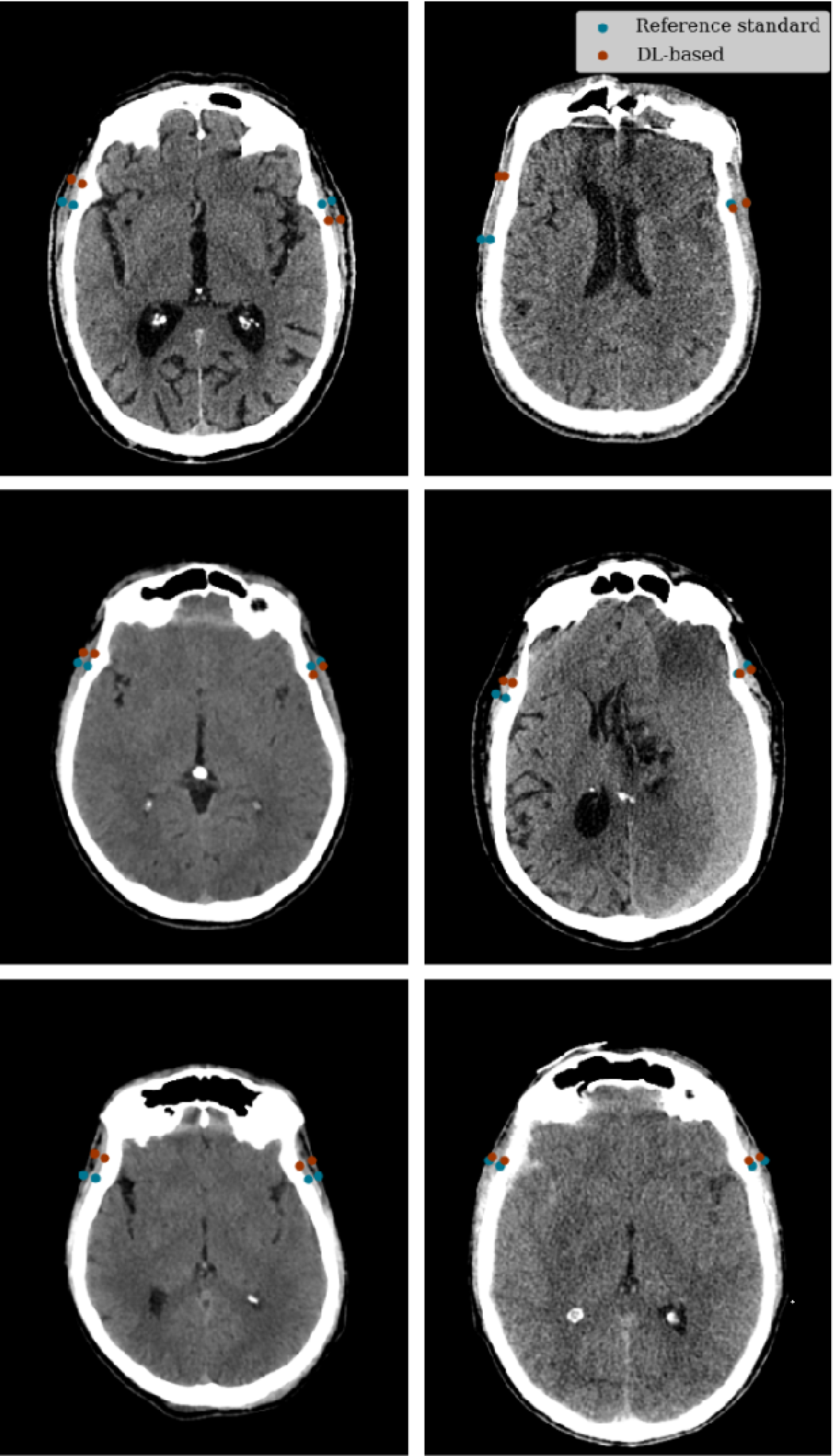
False negative (left) and false positive (right) cases. The TMT measurements of the incorrectly classified cases were within 0.8 mm of the cut-off value. Abbreviations. TMT: temporalis muscle thickness.

Furthermore, the TMT margin (the absolute value of the difference between the DL-based TMT and the cut-off value) of the correctly classified cases had a median of 0.9 mm (IQR: 0.4—1.9), and the margins of the incorrectly classified cases were between <0.1 and 0.8 mm. These findings suggest that the errors and margin of the incorrectly classified cases are not substantially greater than those of the correctly classified cases. The misclassifications can likely be attributed to the fact that the TMT values of the incorrectly classified cases were near the cut-off value, making them highly susceptible to errors.

## Discussion

We have developed and tested a deep learning model for the automatic measurement of temporalis muscle thickness from CT head data. To our knowledge, this is the first study to use DL to quantify TMT from CT head scans. The widespread use and high resolution of modern CT make this a readily accessible modality for the assessment of TMT and, in turn, muscle mass. This makes CT particularly well-suited for scenarios where timely decision-making is critical. The development of automated muscle measurement tools is instrumental in enhancing the recognition of patients with low musculature and thus at-risk of vulnerability. Improved detection of these patients could allow more timely decision-making and direct accurate delivery of ancillary care to those most likely to benefit from it.

This study was limited by the relatively small training set size and the lack of demographic diversity, as the dataset only contained scans collected from institutions within a single city. This could limit the generalizability of our model to broader populations. Nonetheless, the median TMT values measured by the expert clinician in the dataset used were within the ranges reported previously from diverse, international datasets (12), which suggests that the dataset used may be representative of a broader population. To address the sample size limitations, we selected a DL model designed for limited training data, used data augmentation during the training phase to enhance variability, and initialized the model weights with those of a network previously trained on medical data. Finally, this study is limited by the lack of external validation of the model, which was not performed due to a lack of an appropriately annotated external dataset. Furthermore, we could not study the model’s performance with regards to sex and age groups, which are well established confounders of muscle mass, due to the unavailability of this information.

Future work will focus on developing cross-sectional area and volumetric temporalis muscle measurement tools, and comparing their prognostic value to that of TMT, in a larger dataset. Potential barriers for the clinical implementation of these models include variability in acquisition protocol and anatomical differences across patient populations.

In summary, our pipeline has the capacity to accurately estimate TMT and could be used in the recognition of patients with low muscle mass from diagnostic CT head scans. This could be especially useful in acute scenarios where vulnerability of the patient needs to be assessed to inform immediate care, or in the assessment of ancillary care needs.

## Other Information

### Software availability

Scripts for the preprocessing, training, testing, and the weights of the trained model for which results were reported can be found at https://github.com/AngieNicole-Hernandez/TemporalisMuscleSegmentation.

## Bibliography

1. Petermann-Rocha F, Balntzi V, Gray SR, Lara J, Ho FK, Pell JP, et al. Global prevalence of sarcopenia and severe sarcopenia: a systematic review and meta-analysis. J Cachexia Sarcopenia Muscle. 2022 Feb;13(1):86–99.

2. Tolonen A, Pakarinen T, Sassi A, Kyttä J, Cancino W, Rinta-Kiikka I, et al. Methodology, clinical applications, and future directions of body composition analysis using computed tomography (CT) images: A review. Eur J Radiol. 2021 Dec;145:109943.

3. Ten Cate C, Huijs SMH, Willemsen ACH, Pasmans RCOS, Eekers DBP, Zegers CML, et al. Correlation of reduced temporal muscle thickness and systemic muscle loss in newly diagnosed glioblastoma patients. J Neurooncol. 2022 Dec 1;160(3):611–8.

4. Leitner J, Pelster S, Schöpf V, Berghoff AS, Woitek R, Asenbaum U, et al. High correlation of temporal muscle thickness with lumbar skeletal muscle cross-sectional area in patients with brain metastases. PLoS One. 2018;13(11):e0207849.

5. Leitner J, Pelster S, Schöpf V, Berghoff AS, Woitek R, Asenbaum U, et al. High correlation of temporal muscle thickness with lumbar skeletal muscle cross-sectional area in patients with brain metastases. PLoS One. 2018;13(11):e0207849.

6. Park J, Park J, Kim S, Kim DC. Correlation between temporal muscle thickness and grip strength in hemiplegic patients with acute stroke. Front Neurol [Internet]. 2023 Nov 23 [cited 2025 Aug 1];14. Available from: https://www.frontiersin.org/journals/neurology/articles/10.3389/fneur.2023.1252707/full

7. Dubinski D, Won SY, Behmanesh B, Cantré D, Mattes I, Trnovec S, et al. Significance of Temporal Muscle Thickness in Chronic Subdural Hematoma. J Clin Med. 2022 Oct 31;11(21):6456.

8. Korhonen TK, Arponen O, Steinruecke M, Pecorella I, Mee H, Yordanov S, et al. Reduced temporal muscle thickness predicts shorter survival in patients undergoing chronic subdural haematoma drainage. Journal of Cachexia, Sarcopenia and Muscle. 2024;15(4):1441–50.

9. Katsuki M, Kakizawa Y, Nishikawa A, Yamamoto Y, Uchiyama T, Agata M, et al. Temporal Muscle and Stroke-A Narrative Review on Current Meaning and Clinical Applications of Temporal Muscle Thickness, Area, and Volume. Nutrients. 2022 Feb 6;14(3):687.

10. Ravera B, Lombardi C, Bellavia S, Scala I, Cerulli F, Torchia E, et al. Temporalis muscle thickness as a predictor of functional outcome after reperfusion therapies for acute ischemic stroke: a retrospective, cohort study. J Neurol. 2024 Sep 1;271(9):6015–24.

11. Lee B, Bae YJ, Jeong WJ, Kim H, Choi BS, Kim JH. Temporalis muscle thickness as an indicator of sarcopenia predicts progression-free survival in head and neck squamous cell carcinoma. Sci Rep. 2021 Oct 5;11(1):19717.

12. Yang YW, Ming Yang, Zhou YW, Xia X, Jia SL, Zhao YL, et al. Prognostic value of temporal muscle thickness, a novel radiographic marker of sarcopenia, in patients with brain tumor: A systematic review and meta-analysis. Nutrition. 2023 Aug 1;112:112077.

13. Zapaishchykova A, Liu KX, Saraf A, Ye Z, Catalano PJ, Benitez V, et al. Automated temporalis muscle quantification and growth charts for children through adulthood. Nat Commun. 2023 Nov 9;14(1):6863.

14. Huang R, Chen J, Wang H, Wu X, Hu H, Zheng W, et al. Deep learning-based temporal muscle quantification on MRI predicts adverse outcomes in acute ischemic stroke. European Journal of Radiology. 2025 Oct 1;191:112332.

15. Mi E, Mauricaite R, Pakzad-Shahabi L, Chen J, Ho A, Williams M. Deep learning-based quantification of temporalis muscle has prognostic value in patients with glioblastoma. Br J Cancer. 2022 Feb;126(2):196–203.

16. Arponen O, Ikonen JN, Kajantie E, Eriksson JG, Haapanen MJ. Frailty in Late Midlife to Old Age and Its Relationship to Medical Imaging Use and Imaging-related Costs: A Longitudinal Study. Radiology. 2023 Nov;309(2):e230283.

17. Pesonen EK, Arponen O, Niinimäki J, Hernández N, Pikkarainen L, Tetri S, et al. Age- and sex-adjusted CT-based reference values for temporal muscle thickness, cross-sectional area and radiodensity. Sci Rep. 2025 Jan 18;15(1):2393.

18. Chilamkurthy S, Ghosh R, Tanamala S, Biviji M, Campeau NG, Venugopal VK, et al. Development and Validation of Deep Learning Algorithms for Detection of Critical Findings in Head CT Scans [Internet]. arXiv; 2018 [cited 2025 Jul 28]. Available from: http://arxiv.org/abs/1803.05854

19. Jonkers J, Duchateau L, Van Wallendael G, Van Hoecke S. landmarker: A Toolkit for Anatomical Landmark Localization in 2D/3D Images. SoftwareX. 2025 May 1;30:102165.

20. Payer C, Štern D, Bischof H, Urschler M. Integrating spatial configuration into heatmap regression based CNNs for landmark localization. Medical Image Analysis. 2019 May 1;54:207–19.

21. Cardoso MJ, Li W, Brown R, Ma N, Kerfoot E, Wang Y, et al. MONAI: An open-source framework for deep learning in healthcare [Internet]. arXiv; 2022 [cited 2025 Jul 28]. Available from: http://arxiv.org/abs/2211.02701

22. Akiba T, Sano S, Yanase T, Ohta T, Koyama M. Optuna: A Next-generation Hyperparameter Optimization Framework. In: Proceedings of the 25th ACM SIGKDD International Conference on Knowledge Discovery & Data Mining [Internet]. New York, NY, USA: Association for Computing Machinery; 2019 [cited 2025 Aug 16]. p. 2623–31. (KDD ’19). Available from: 10.1145/3292500.3330701

23. Dubinski D, Won SY, Meyer-Wilmes J, Trnovec S, Rafaelian A, Behmanesh B, et al. Frailty in Traumatic Brain Injury—The Significance of Temporal Muscle Thickness. Journal of Clinical Medicine. 2023 Jan;12(24):7625.

